# Maternal microbes and early brain development in mouse

**DOI:** 10.1101/2022.12.19.521137

**Authors:** Xin Yi Yeo, Woo Ri Chae, Hae Ung Lee, Han-Gyu Bae, Sven Pettersson, Joanes Grandjean, Weiping Han, Sangyong Jung

**Affiliations:** Institute of Molecular and Cell Biology, Agency for Science, Technology and Research, Singapore; Department of Psychological Medicine, Yong Loo Lin School of Medicine, National University of Singapore, Singapore; Department of BioNano Technology, Gachon University, Seongnam, The Republic of Korea; National Neuroscience Institute, Tan Tock Seng Hospital, Singapore Health Services, Singapore; University of Texas Health Science Center at San Antonio, United States; Department of Microbiology and Immunology, Yong Loo Lin School of Medicine, National University of Singapore, Singapore; Department of Medical Sciences, Sunway University, Kuala Lumpur, Malaysia; Department of Medical Imaging, Radboud University Medical Center, Nijmegen, The Netherlands; Donders Institute for Brain, Cognition and Behaviour, Radboud University Medical Center, Nijmegen, The Netherlands; Department of Physiology, Yong Loo Lin School of Medicine, National University of Singapore, Singapore

**Keywords:** Germ-free Mice, Gut Microbiome, Early-life Alteration, Brain Development, MRI

## Abstract

The complex symbiotic relationship between the mammalian body and gut microbiome plays a critical role in the health outcomes of offspring later in life. The gut microbiome modulates virtually all physiological functions through direct or indirect interactions to maintain physiological homeostasis. Previous studies indicate a link between maternal/early-life gut microbiome, brain development, and behavioral outcomes relating to social cognition. Here we present direct evidence of the role of the gut microbiome in brain development. Through magnetic resonance imaging (MRI), we investigated the impact of the gut microbiome on brain organization and structure using germ-free (GF) mice and conventionalized mice, with the gut microbiome reintroduced after weaning. We found broad changes in brain volume in GF mice that persist despite the reintroduction of gut microbes at weaning. These data suggest a direct link between the maternal gut or early-postnatal microbe and their impact on brain developmental programming.

## 1. Introduction

The mammalian body is inhabited by a myriad of microorganisms originating from the vertical transfer of vaginal-perianal microbes during delivery (1,2). The complex symbiotic relationship rapidly established after birth between the offspring and the microorganisms hosted (3), particularly within the gastrointestinal system, is critical for the establishment and maturation of immune functions (4), metabolism (5,6), and the modulation of physiological homeostasis (7,8). Through direct anatomical connection, indirect endocrine means, or the mobilization of the immune system, microbial metabolites from the GI tract can modulate a broad range of physiological functions (9,10).

It is increasingly apparent that the gut microbiota is a direct, modifiable factor of neurodevelopment and neurological outcomes in humans. Early exposure to antibiotics, which alters microbiome diversity (11,12), is associated with an increased risk of cognitive and neurodevelopmental disorders such as attention-deficit hyperactivity disorder and autism spectrum disorder (13,14). Fecal transplantation performed on germ-free (GF) dams using human maternal microbes from preterm mothers suggests poor growth outcomes (15). Interestingly, there is a correlation between the time and duration of antibiotic exposure and the severity of cognitive deficits later in life (16), and the levels of gut microbe by-products 3,4-dihydroxyphenylacetic acid and sodium butyrate are positively associated with mental health (17,18) and adult neurogenesis (19).

Experiments using the offspring of germ-free GF and antibiotic-treated dams further revealed that planar indoles and amine oxides produced by gut microbes regulate the expression of genes involved in neuronal differentiation and axonogenesis, directly impacting neurogenesis (20,21). In relation, Sven Pettersson and others observed that altered gut microbiome colonization and composition significantly impact cognitive behaviour in mice (22–26), with a preferential effect on social behaviour. Although pieces of information point to the involvement of gut microbiota in the modulation of brain development and function (27), there is a lack of direct evidence of gut microbiome-dependent change in brain development.

Preclinical magnetic resonance imaging (MRI) methods provide a unique opportunity for a complete brain structure examination under the defined absence of gut microbiome. In this study, we examined the impact of early life absence of gut microbiome on brain development through structural MRI between control, specific-pathogen-free mice carrying normal microbiome population and the GF mice (Figure 4). Alterations in isocortex and olfactory cortex volumes associated with higher-order cognitive control ensued from the complete depletion of microorganisms sustained through a restriction in environmental exposure. Additionally, the conventionalization of young adult GF mice failed to normalize the brain changes observed. These observations provide a premise for future functional and neurocircuitry studies to explore the structural determinants of cognitive deficits associated with early-life disturbance in the gut microbiome.

## 2. Materials and Methods

### 2.1 Animal welfare

All animal procedures were carried out per institutional guidelines by Nanyang Technological University and approved by Regional Animal Research Ethical Board, Institutional Animal Care and Use Committee, Singapore. Male NMRI mice, 6-8 weeks old, are used for MRI analysis in this study. All mice were kept under a 12-h light/dark cycle and provided with water and food (autoclaved R36 Lactamin) ad libitum. GF mice were placed and raised in special sterile plastic isolators. The isolator sterility was analyzed weekly through the bacterial and fungal culture of fecal homogenates of GF mice. Part of the GF mice population was conventionalized (CONV) after weaning (between 3 to 4 weeks of age) by gavage of 100 μl of specific pathogen-free (SPF) mouse fecal homogenate (dissolving two fecal pellets from SPF mice in 1 ml of phosphate-buffered saline) and cohoused with SPF mice for 21 days before being euthanized. 5 mice from each group are used in this study. No subjects were excluded from the analysis.

### 2.2 MRI acquisition and data processing

Mouse brains are imaged postmortem *in situ* (28). Data were acquired with a Bruker 11/16 Biospec spectrometer (Bruker BioSpin MRI, Ettlingen, Germany) operating at 11.75 T, equipped with a BGA-S gradient system, a linear volume resonator coil for transmission, and a two-by-two phased-array cryocoil surface receiver coil. Images were acquired using Paravision 6 software. Localizer images were acquired to ensure the correct positioning of mice to the coil and the magnet isocentre. 3D volumetric scanning occurred for 8h30 min, with scans acquired using spin echo RARE sequence (29) with a field of view saturation slice positioned on the inferior portion of the head and acquisition parameters: repetition time = 2500 ms, echo time = 5.1 ms, RARE factor = 10, matrix size 250 × 225 × 250, and field of view =20 × 18 × 20 mm^3^, average = 2, achieving a resolution of 0.08 mm^3^.

Anatomical scans were processed through voxel-based morphometry using ANTS (Advanced Normalization Tools 2.4.0, picsl.upenn.edu/software/ants). Briefly, images were first corrected for intensity field inhomogeneity using the N4 algorithm. Second, a study template was estimated by realigning individual volumes to each other over 3 iterative steps of non-linear transformation. Third, the individual volumes underwent SyN diffeomorphic image registration to the study template. Fourth, the Jacobian determinants were estimated and voxel-wise comparisons between groups were carried out with a two-sample t-test using FSL 6.0.5 (fsl_glm) and corrected for multiple hypothesis testing using the easy threshold function.

### 2.3 Data availability

The raw datasets generated for this study are available in http://doi:10.18112/openneuro.ds004254.v1.0.0. The programming code used for data preprocessing and analysis can be found at https://gitlab.socsci.ru.nl/preclinical-neuroimaging/germfree. Further inquiries can be directed to the corresponding authors.

### 2.4 Statistical analysis

Jacobian determinants were extracted using the DSURQE mouse brain atlas (30). All data are expressed as mean ± standard error mean (SEM) and ported to GraphPad Prism 7.04 (GraphPad Software) for analysis. Datasets were tested for normal distribution with the Shapiro-Wilk test. Further analysis was conducted with the Mann-Whitney U test between SPF, GF, and CONV datasets. P values < 0.05 were considered statistically significant.

## 3. Results

### 3.1 GF mice exhibit distinct brain volume changes compared with age-matched SPF mice

Associations between early-life microbiome dysbiosis and later-stage cognitive deficits suggest neurodevelopmental deviations under gnotobiotic conditions (16,31,32). Yet the loci and extent of neurological changes are unknown. To identify brain regions modulated by commensal bacteria, volumetric analysis of 6 to 8 weeks old SPF and GF mouse brains was achieved with MRI. Corrected statistical maps depicting significant group differences (p corrected < 0.05) are shown with an anatomical underlay. We observed a widespread alteration in brain volume in the GF versus the SPF mouse brain (Figure 1A). Notably, the absence of microbiome results in a significant bilateral increase in the olfactory cortex and isocortical brain volumes (AP +1.98 to -3.28, orange-red sections) and a decrease in the sensory-related superior colliculus (SCs; AP -4.96), thalamus (TH, AP -2.8), and hypothalamus (AP -0.94; blue-green sections) in the GF compared with SPF mouse (Figure 1A). Specifically, there is a significant increase in the volumes of the somatomotor (Total MO, includes MOp and MOs, 0.044 ± 0.006, p = 0.016), somatosensory (SSp, includes SSp-bfd, SSp-ul, SSp-ll, SSp-m, SSp-tr and SSp-n,, 0.073 ± 0.014, p = 0.016; SSs 0.073 ± 0.010, p = 0.008), anterior cingulate cortex (ACA, including ACAd and ACAv, p = 0.016), prelimbic area (PL 0.105 ± 0.026, p = 0.032), ectorhinal area (ECT 0.068 ± 0.009, p = 0.008), frontal pole of cerebral cortex (FRP 0.074 ± 0.010, p = 0.008), temporal association areas (TEa 0.063 ± 0.017, p = 0.016) and the perihinal area (PERI 0.072 ± 0.014, p = 0.016) of the isocortex, and the anterior olfactory nucleus (AON 0.094 ± 0.017, p = 0.032) and taenia tecta (TT, includes TTv and TTd, 0.101 ± 0.015, p = 0.008) of the olfactory cortex in GF mice (Figure 1B and Supplementary Figure 1).

**Figure 1:**
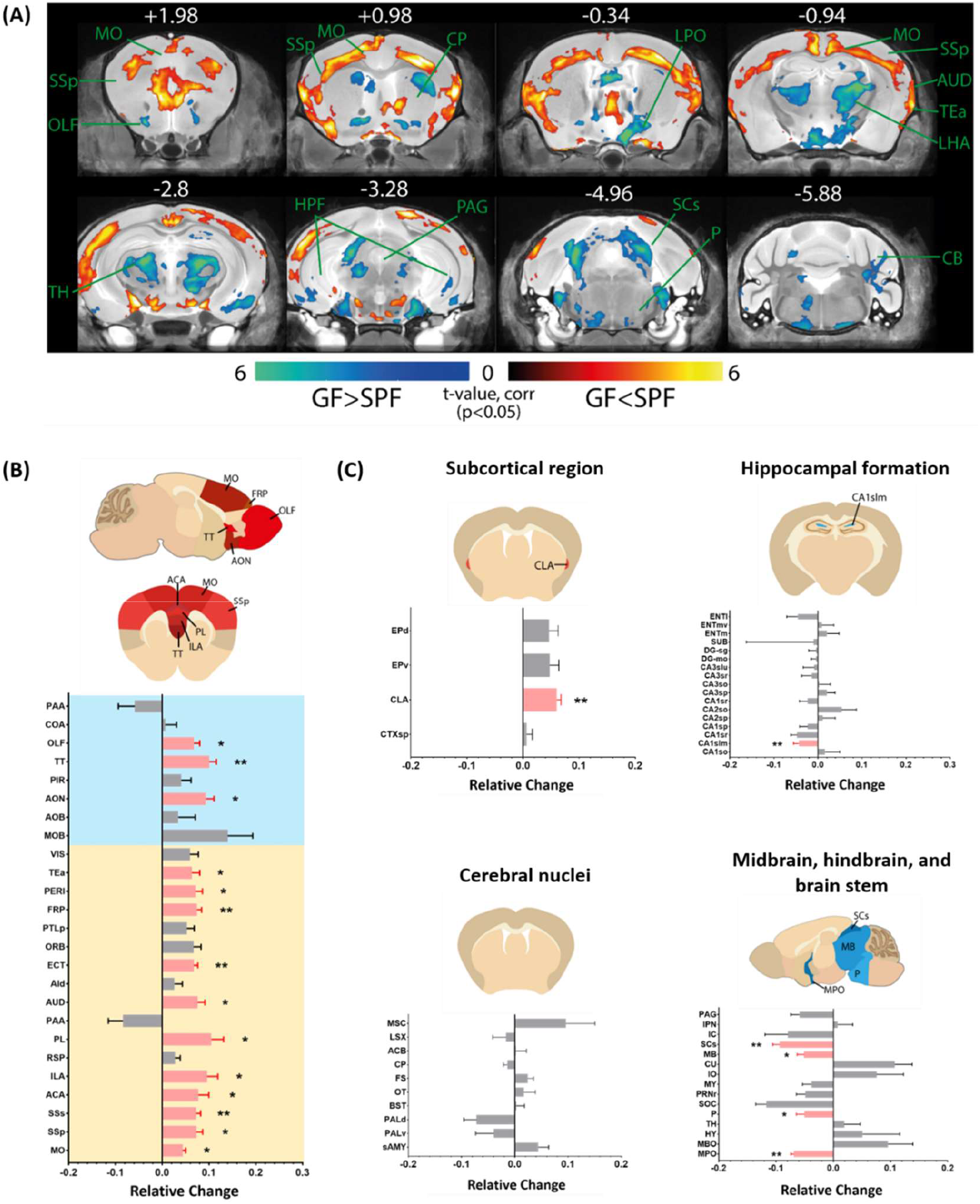
Widespread alterations in the subcortical brain volume of germ free (GF) mice. (A) Coronal view of brain regions exhibiting reduced (blue-green) or increased (yellow-red) brain volume in GF compared to age-matched SPF subjects (control). The AP coordinates (in mm) of the exact coronal sections location in the mouse brain are provided on top of each of the presented statistical maps. MO somatomotor cortex, SSp primary somatosensory cortex, OLF olfactory cortex, CP caudoputamen, LPO lateral preoptic area, AUD auditory area, TEa temporal association area, LHA lateral hypothalamic area, TH thalamus, HPF hippocampal formation, PAG periaqueductal gray, SCs sensory-related superior colliculus, P pons, CB cerebellum. (B) Relative brain volume changes of age-matched SPF compared with GF, normalized against the average brain volume from the SPF subjects. Orange, isocortex; Blue, olfactory cortex. (C) Relative subcortical region, hippocampal formation, cerebral nuclei, and the midbrain, hindbrain, and brain stem volume changes of age-matched SPF compared with GF normalized against the average brain volume from the SPF subjects. The precise names of brain regions can be found in the list of abbreviations in supplementary materials. Brain regions are arranged based on their anatomical location. SPF n = 5, GF n = 5. Data presented as mean ± SEM. *, p < 0.05; **, p < 0.01.

### 3.2 Conventionalization post-weaning does not reverse major brain volume changes observed in GF mice

To further understand the specific brain volume differences observed between the SPF and GF mice and examine the impact of the absence of gut microbiome on brain volume changes, we reviewed the effect of reintroducing gut microbiome post-weaning (conventionalization) on brain volumes of GF mice (CONV) compared against age-matched SPF mice (Figure 2A). Despite having a fetal bacterial population resembling the donor SPF mice post conventionalization (33), brain areas altered in conventionalized GF mice are similar to that of GF mice compared against the SPF mice (Figure 2B; MO 0.054 ± 0.008, p = 0.016; SSp 0.069 ± 0.006, p = 0.008; SSs 0.052 ± 0.015, p = 0.008; ACA 0.090 ± 0.011, p = 0.008; PL 0.145 ±0.032, p = 0.008; ECT 0.064 ± 0.006; p = 0.008; FRP 0.116 ± 0.025, p = 0.032; TEa 0.044 ± 0.011, p = 0.032; PERI 0.079 ± 0.004, p = 0.008; AON 0.125 ± 0.011, p = 0.008; TT 0.107 ± 0.017, p = 0.008, all values in 3 d.p.), and that volumes of these regions are comparable between CONV and GF mice (Figure 3B, Supplementary Figure 1; MO 0.009 ± 0.008, p = 0.421; SSp -0.004 ± 0.006, p > 0.999; SSs -0.020 ± 0.014, p = 0.421; ACA 0.011 ± 0.010, p > 0.999; PL 0 ± 0.010, p = 0.548; ECT 0 ± 0.008, p = 0.691; FRP 0 ± 0.009; p = 0.151; TEa -0.018 ± 0.010, p = 0.421; PERI 0 ± 0.013, p = 0.841; AON 0.028 ± 0.011, p = 0.222; TT 0.005 ± 0.016, p > 0.999, all values in 3 d.p).

**Figure 2:**
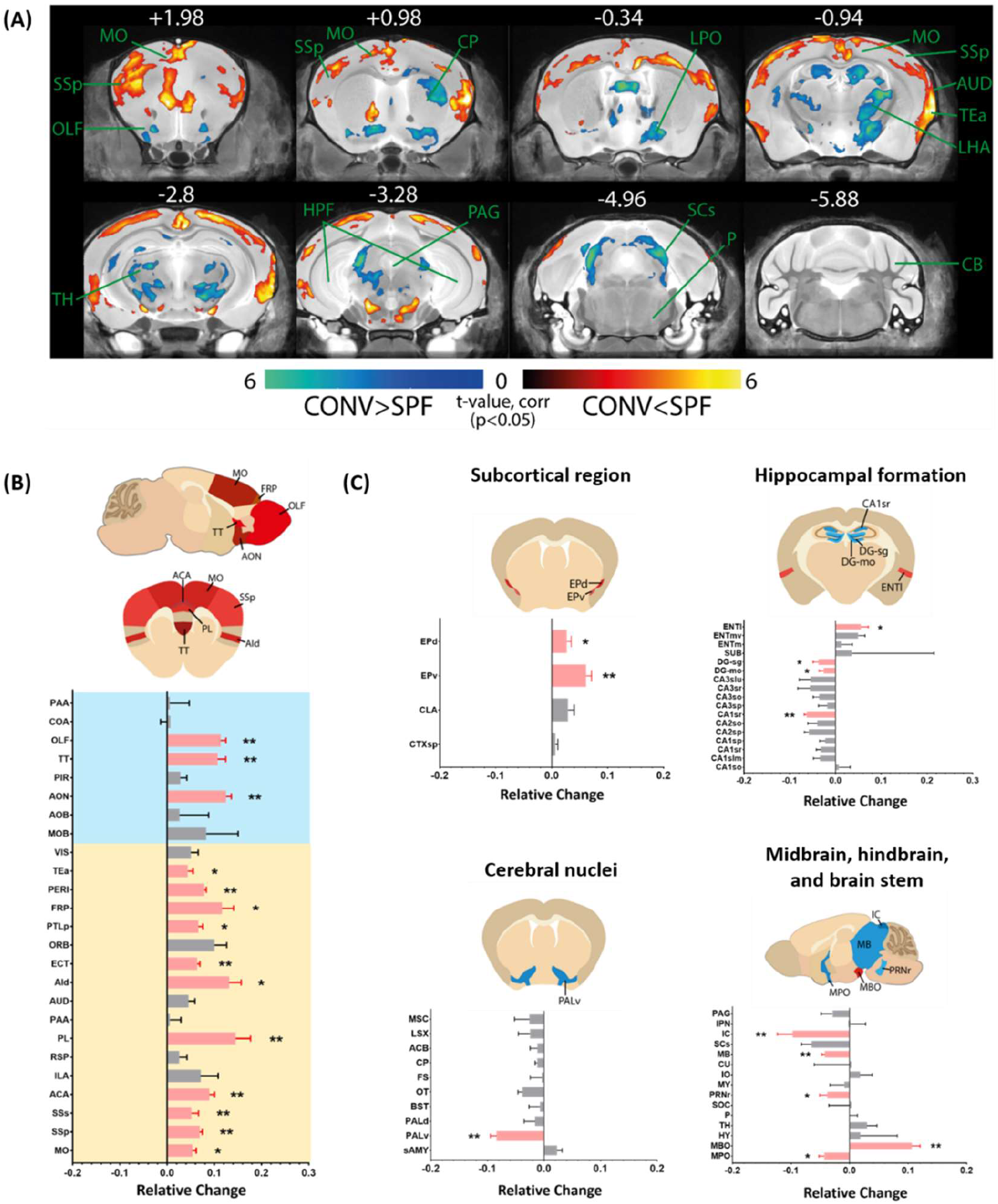
Conventionalisation of GF mice post-weaning does not reverse majority of the brain volume changes in GF mice. (A) Coronal view of brain regions exhibiting reduced (blue-green) or increased (yellow-red) brain volume in post-weaning conventionalised GF mice (CONV) compared to age-matched SPF subjects (control). The AP coordinates (in mm) of the exact coronal sections location in the mouse brain are provided on top of each of the presented statistical maps. MO somatomotor cortex, SSp primary somatosensory cortex, OLF olfactory cortex, CP caudoputamen, LPO lateral preoptic area, AUD auditory area, TEa temporal association area, LHA lateral hypothalamic area, TH thalamus, HPF hippocampal formation, PAG periaqueductal gray, SCs sensory-related superior colliculus, P pons, CB cerebellum. (B) Relative brain volume changes of age-matched SPF compared with CONV, normalized against the average brain volume from the SPF subjects. Orange, isocortex; Blue, olfactory cortex. (C) Relative subcortical region, hippocampal formation, cerebral nuclei, and the midbrain, hindbrain, and brain stem volume changes of age-matched SPF compared with CONV normalized against the average brain volume from the SPF subjects. The precise names of brain regions can be found in the list of abbreviations in supplementary materials. Brain regions are arranged based on their anatomical location. SPF n = 5, CONV n = 5. Data presented as mean ± SEM. *, p < 0.05; **, p < 0.01.

**Figure 3:**
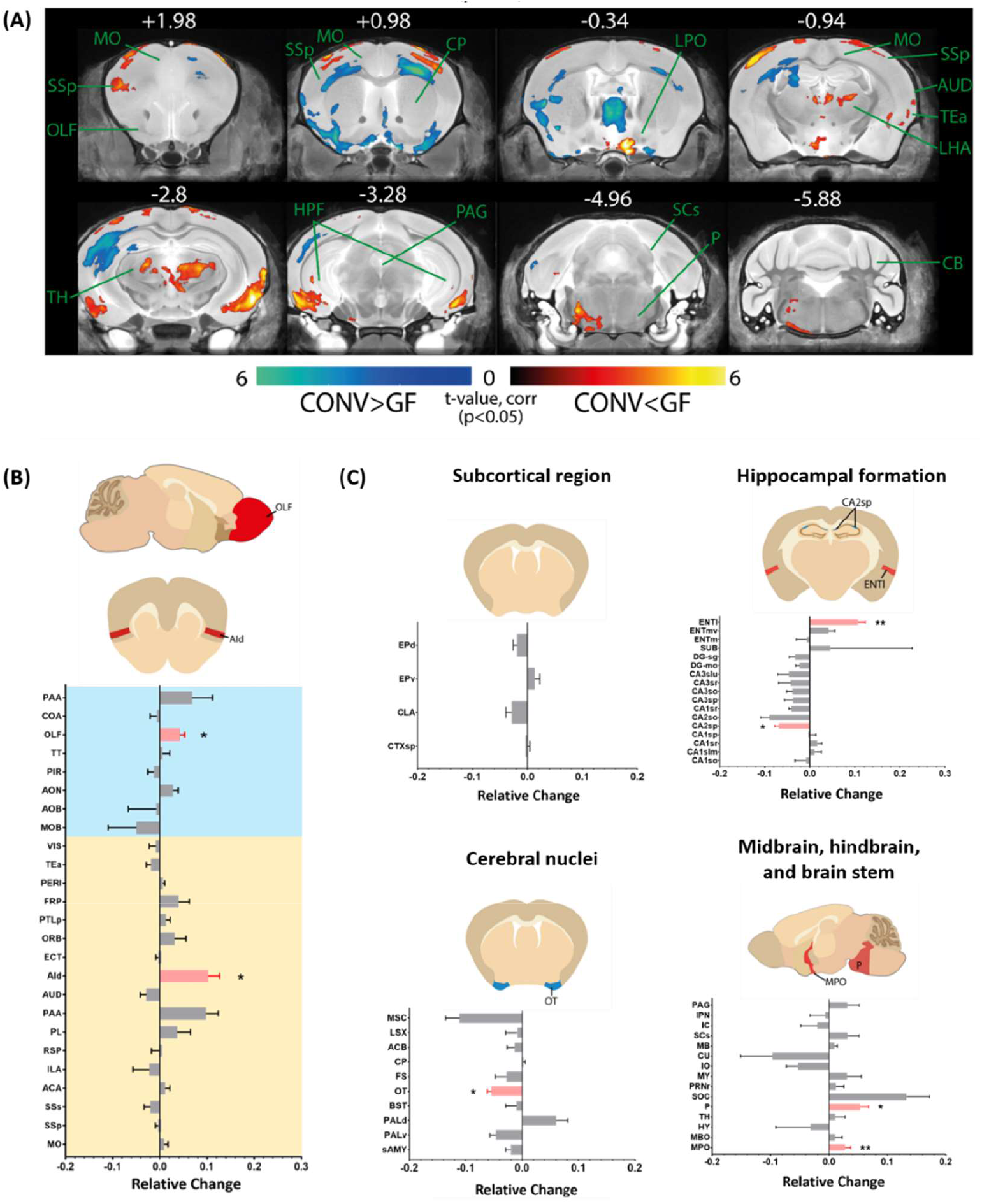
Brain regions sensitive toward physiological conditions are affected by the reintroduction of gut microbiome. (A) Coronal view of brain regions exhibiting reduced (blue-green) or increased (yellow-red) brain volume in GF compared to age-matched post-weaning conventionalised GF mice (CONV) subjects. The AP coordinates (in mm) of the exact coronal sections location in the mouse brain are provided on top of each of the presented statistical maps. MO somatomotor cortex, SSp primary somatosensory cortex, OLF olfactory cortex, CP caudoputamen, LPO lateral preoptic area, AUD auditory area, TEa temporal association area, LHA lateral hypothalamic area, TH thalamus, HPF hippocampal formation, PAG periaqueductal gray, SCs sensory-related superior colliculus, P pons, CB cerebellum. (B) Relative brain volume changes of age-matched GF compared with CONV, normalized against the average brain volume from the SPF subjects. Orange, isocortex; Blue, olfactory cortex. (C) Relative subcortical region, hippocampal formation, cerebral nuclei, and the midbrain, hindbrain, and brain stem volume changes of age-matched GF compared with CONV normalized against the average brain volume from the SPF subjects. The precise names of brain regions can be found in the list of abbreviations in supplementary materials. Brain regions are arranged based on their anatomical location. SPF n = 5, CONV n = 5. Data presented as mean ± SEM. *, p < 0.05; **, p < 0.01.

### 3.3 The reintroduction of gut microbiome influenced brain regions sensitive towards general physiological conditions

Gut microbiome disturbances can induce permanent alterations in key neurological processes and brain development or transient, reversible metabolic-dependent fluctuations in brain volume. The reintroduction of microorganisms postnatally is sufficient to normalise the subcortical claustrum (CLA SPF vs GF 0.060 ± 0.008, p = 0.008; SPF vs CONV 0.029 ± 0.011, p = 0.151), the isocortex infralimbic area (ILA SPF vs GF 0.096 ± 0.023, p = 0.016; SPF vs CONV 0.071 ± 0.04, p = 0.222), auditory area (AUD SPF vs GF 0.076 ± 0.016, p = 0.032; SPF vs CONV 0.045 ± 0.014, p = 0.095), hippocampal formation CA1 region (CA1slm SPF vs GF -0.043 ± 0.012, p = 0.008; SPF vs CONV 0.010 ± 0.015, p = 0.056), the pons (P SPF vs GF -0.084 ± 0.014, p = 0.016; SPF vs CONV -0.009 ± 0.020, p = 0.548), and the brain stem superior colliculus (SCs SPF vs GF -0.093 ± 0.012, p = 0.008; SPF vs CONV -0.065 ± 0.018, p = 0.056) (Figure 2B and C). Surprisingly, we observed a significant alteration in the thalamic nuclei (AP-0.34 to -2.8) and pons (AP -4.96) of the CONV mouse versus GF mouse brain (Figure 3A) and alterations in previous unaffected dorsal agranular insular area (Figure 1B, 2B and 3B; Aid), posterior parietal association area (PTLp), and lateral entorhinal area (Figure 1C, 2C and 3C; ENTl), endopiriform nucleus (EP consisting of EPd and EPv), dentate gyrus (DG), and the ventral pallidum (PALv). The fiber tract and ventricular system are unaffected by the altered gut microbiome composition (Supplementary Figure 2).

## 4. Discussion

Epidemiological studies have reported a possible correlation between alterations in the composition of the gut microbiome and function and increased risk of neurodevelopmental and neuropsychiatric disorders (13,34). Preclinical animal studies further showed that exposure to gut microbial products can alter key neurodevelopmental processes (20,21,35,36). However, the direct contribution of the gut microbiota and neurocognitive outcomes remains inconclusive due to a lack of understanding of the underlying mechanisms (37–39). Here, we present evidence for an association between structural alterations in the olfactory cortex and isocortex of GF mice (Figure 4) and the maternal microbe’s regulation of developmentally regulated structural changes associated with the growth of offspring in utero.

**Figure 4:**
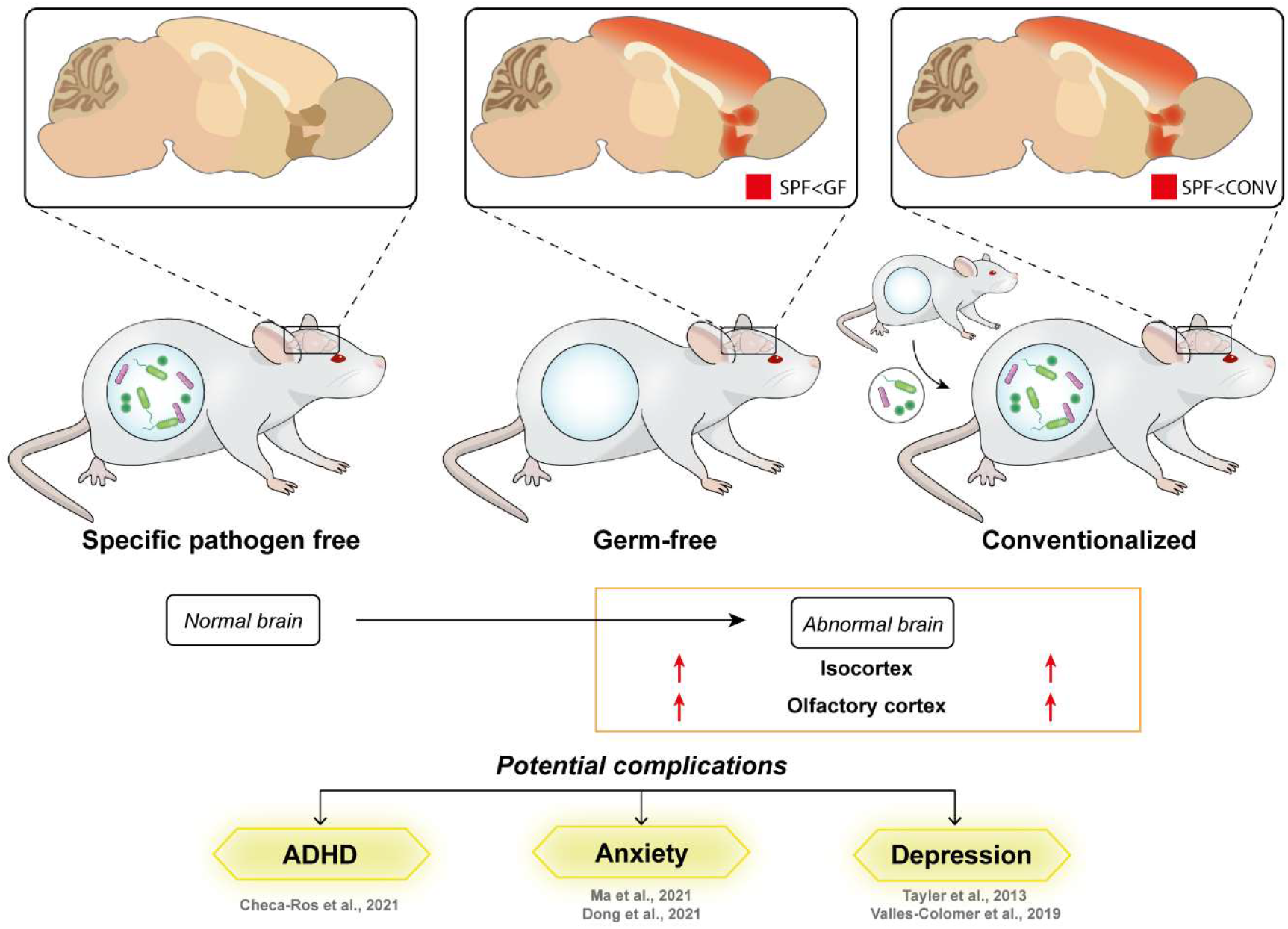
Ablation of gut microbiota persistently affects brain development in young adult mice. An increase in the volume of olfactory and somatosensory cortices observed in the absence of microorganisms in the GF mice is irreversible with the reintroduction of the normal gut microbiome from the SPF mice in early postnatal life. As such, the resulting brain volume changes in CONV mice are similar to that of GF mice compared to SPF mice. Alterations in development and the neuronal circuitry of these brain regions are often associated with the development of neuropsychiatric conditions such as attention deficit hyperactivity disorder (ADHD), anxiety, and depression.

The increase of interest in the effects of intestinal microbiota on human health has led to extensive metagenome sequencing efforts to understand the variation and diversity between organisms and health conditions (40–43) and the generation of preclinical models to elucidate the gut microbiome mechanism of action (44–46). The use of GF mice as the basis of the bidirectional gut microbiome-brain communication provides the opportunity to decipher the role of the gut microbiota and regulation of brain development and function.

Brain volume change is a powerful correlate of neurodevelopmental and neurocognitive outcomes of individuals (47,48), and MRI is a non-invasive method that allows for the complete examination of brain structure changes in a single experimental run. We observed broad structural volume alteration specific to the somatosensory in GF mice, critical for general and higher-order sensory processing and integration (49–53), compared to their age-matched SPF counterparts (Figure 1B, 3B, and Supplementary Figure 1). This is consistent with previous reports detailing defects in sensory processing in neuropsychiatric conditions associated with a disturbed gut microbiome (13,54,55). In addition, changes in the isocortex infralimbic area (ILA) and anterior cingulate area (ACA) associated with contextual fear conditioning (56) and the modulation of fear and anxiety (57–59) correspond to altered anxiety behaviour commonly observed in GF mice (22,23,60).

Furthermore, there is a directed increase in the volumes of the anterior olfactory nucleus (AON) and taenia tecta (TT) of the olfactory cortex (Figure 1B, 3B, and Supplementary Figure 1), which function as key areas for olfactory information processing and integration (61). Although relatively less explored, olfactory compromises are often observed in autism spectrum disorder patients (62,63) and correlated with the development of depression (64) and schizophrenia (65). The olfactory nerve undergoes a distinct developmental process (66), thought to involve the function of yet unknown odorant receptors (67), which may be directly responsive to gut microbiome-produced metabolites such as short-chain fatty acids and volatile amines that functions as odorants (68,69). A deep examination of the interactions between gut microbiome activities and the olfactory system is required to understand the relevance of microbiome disturbance in the modulation of olfactory development and behaviour.

Gut microbe metabolites were shown to regulate the expression of genes and processes involved in neural stem cell differentiation, neurogenesis, and development (20,21,70). Also, gut-derived neuroactive substances such as tryptophan and serotonin modulate the inflammatory status and function of the microglia in the central nervous system (CNS) (71,72). Due to the time-sensitivity of the production of morphogens (73), and the vulnerability of the neurological system to changes during key neurodevelopmental stages, neurological deviations can lead to persistent structural, functional, or cognitive disability. In rodents, key neurodevelopmental milestones are concluded by a month of age (postnatal) (74), corresponding to the timepoint of gut microbiome reintroduction to the GF mice in this study. Notably, reintroducing the gut microbiome through conventionalization at 3 to 4 weeks of age cannot reverse the majority of observed brain alterations (Figure 2B and C), in line with a previous report suggesting that behavioural changes in NMRI mice can only be reversed with postnatal colonization of microbes from females transplanted with microbes before pregnancy (22). The conventionalization of GF mice is nonetheless sufficient to evoke changes in the colliculus superior (SCs) and hippocampus (Supplementary Figure 2), reflecting their sensitivity toward metabolic alterations (75–77).

We recognize that brain volume changes are dynamic and can be affected by short-term deviations in physiological conditions (78). Furthermore, it is not possible to distinguish the role of maternal gestational microbiota and early postnatal microbiome in brain volume changes observed, and the brain volume alterations do not reveal the underlying structural vulnerability responsible for the cognitive correlates observed in individuals with gut microbiome disturbance. It is unclear how the difference in weight and body fat content of GF mice from an altered gut microbiome-dependent metabolic profile may contribute to changes in brain development (79). Treatment of neurodevelopmental disorders in later stages of life is a possible concept for the normalization of neuronal network function without impacting the underlying defect in consideration of adult neurological plasticity. Further work is required to investigate the mechanisms involved in the observations made in the current report.

## 6. Acknowledgements

This work is supported by the Joint Council Office grant (BMSI/15-800003-SBIC-00E) from Agency for Science, Technology and Research (A*STAR), Singapore. S.P. is supported by the ASEAN Microbiome Nutrition Centre (AMNC), Singapore, UK Dementia group, and National Neuroscience Institute (NNI), SingHealth, Singapore.

## 7. Author Contributions

Conceptualisation, X.Y.Y., S.P., and S.J.; data acquisition and analysis, J.G., X.Y.Y., and H.U.L.; writing – original draft preparation. X.Y.Y., W.R.C., and S.J.; writing – review and editing, X.Y.Y., S.P., J.G., W.H., and S.J.; supervision, S.J.; funding acquisition S.P., W.H., and S.J.; critical revision of manuscript, S.J., S.P., W.H., and J.G. All authors have read and agreed to the published version of the manuscript.

## Supplementary materials for

**Supplementary Figure 1:**
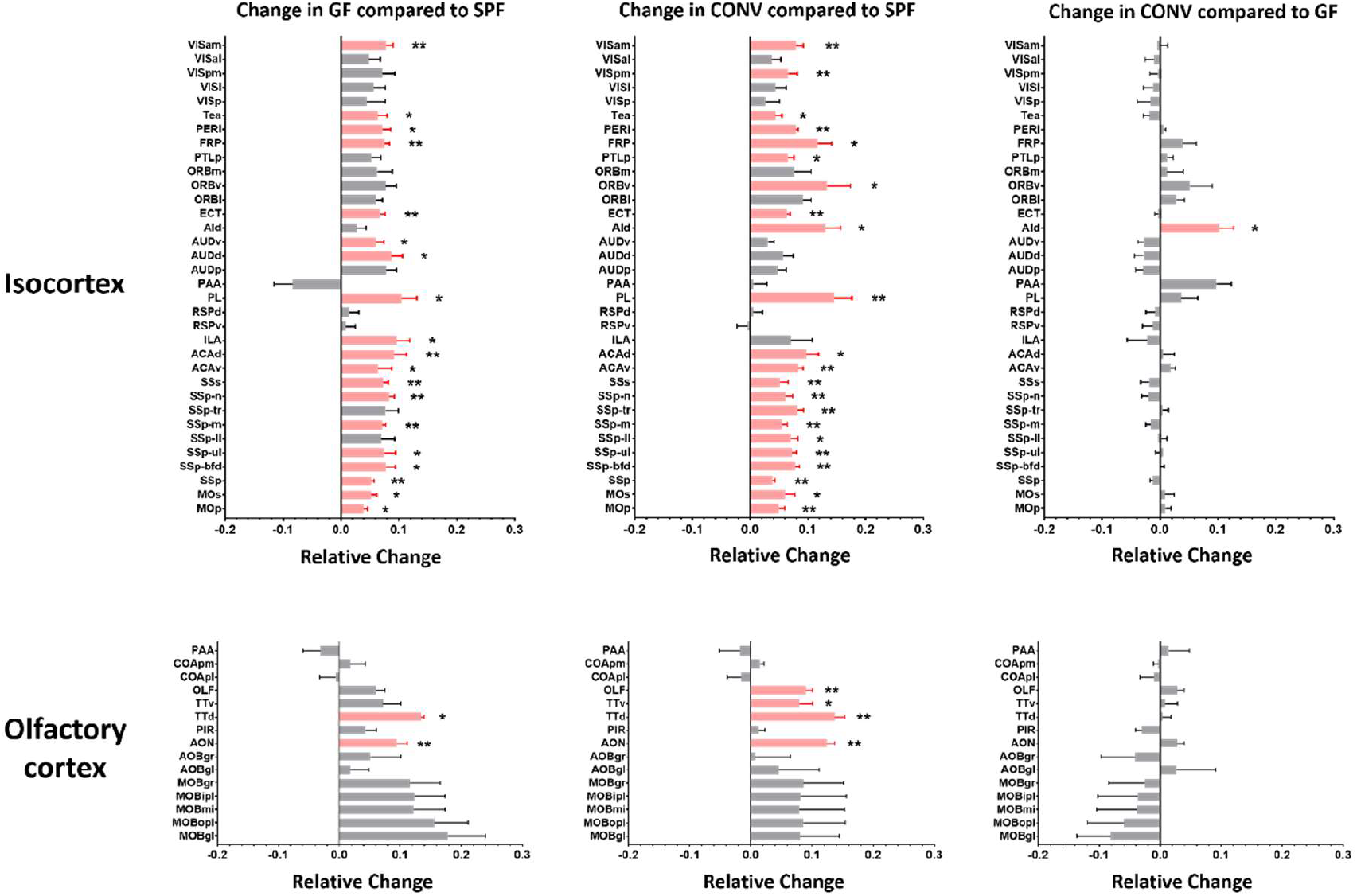
Relative change in volume of defined regions of the isocortex and olfactory cortex of the specific pathogen free (SPF), germ-free (GF) and conventionalized (CONV) mouse brain. The full name of each of the brain region listed can be found in the table of abbreviations. Brain areas with a significant deviation (p < 0.05) are presented in red. Brain areas with a significant deviation (p < 0.05) are presented in red. Data presented as mean ± SEM. *, p < 0.05; **, p < 0.01.

**Supplementary Figure 2:**
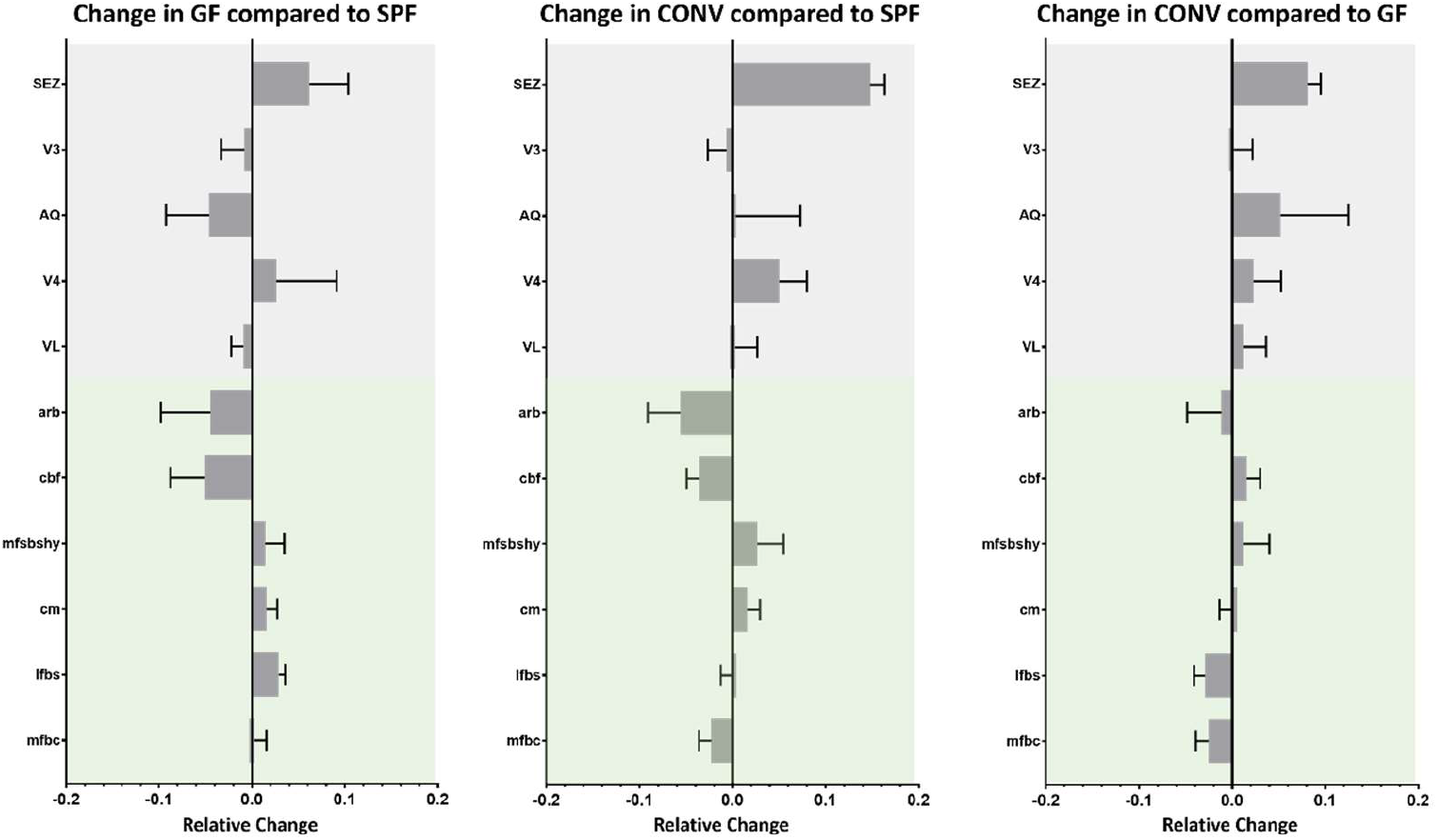
Relative change in volume of the major fibre tracts and the ventricular system of the specific pathogen free (SPF), germ-free (GF) and conventionalized (CONV) mouse brain. Green Fiber tracts; Gray Ventricular system. The full name of each of the brain region listed can be found in the table of abbreviations. Data presented as mean ± SEM.

## Abbreviations

CTXsp: Subcortical region, cortical subplate
CLA: Subcortical region, claustrum
EP: Subcortical region, endopiriform nucleus
EPd: Subcortical region, endopiriform nucleus, dorsal part
EPv: Subcortical region, endopiriform nucleus, ventral part
MO: Isocortex, somatomotor areas
MOp: Isocortex, somatomotor areas, primary motor area
MOs: Isocortex, somatomotor areas, secondary motor area
SSp: Isocortex, primary somatosensory area
SSp.bfd: Isocortex, primary somatosensory area, barrel field
SSp.ul: Isocortex, primary somatosensory area, upper limb
SSp.II: Isocortex, primary somatosensory area, lower limb
SSp.m: Isocortex, primary somatosensory area, mouth
SSp.tr: Isocortex, primary somatosensory area, trunk
SSp.n: Isocortex, primary somatosensory area, nose
SSs: Isocortex, supplemental primary somatosensory area
ACA: Isocortex, anterior cingulate area
ACAv: Isocortex, anterior cingulate area, ventral part
ACAd: Isocortex, anterior cingulate area, dorsal part
ILA: Isocortex, infralimbic area
RSP: Isocortex, retrosplenial area
RSPv: Isocortex, retrosplenial area, ventral part
RSPd: Isocortex, retrosplenial area, dorsal part
PL: Isocortex, prelimbic area
PAA: Isocortex, postpiriform transition area
AUD: Isocortex, auditory area
AUDp: Isocortex, auditory area, primary auditory area
AUDd: Isocortex, auditory area, dorsal auditory area
AUDv: Isocortex, auditory area, ventral auditory area
AId: Isocortex, agranular insular area, dorsal part
ECT: Isocortex, ectorhinal area
ORB: Isocortex, orbital area
ORBl: Isocortex, orbital area, lateral part
ORBv: Isocortex, orbital area, ventrolateral part
ORBm: Isocortex, orbital area, medial part
PTLp: Isocortex, posterior parietal association areas
FRP: Isocortex, frontal pole, cerebral cortex
PERI: Isocortex, perihinal area
TEa: Isocortex, temporal association areas
VIS: Isocortex, visual area
VISp: Isocortex, visual area, primary visual area
VISl: Isocortex, visual area, lateral visual area
VISpm: Isocortex, visual area, posteromedial visual area
VISal: Isocortex, visual area, anterolateral visual area
VISam: Isocortex, visual area, anteromedial visual area
CA1: Hippocampal formation, Ammon’s horn, Field CA1
CA1so: Hippocampal formation, Ammon’s horn, Field CA1, stratum oriens
CA1slm: Hippocampal formation, Ammon’s horn, Field CA1, stratum lacunosum-moleculare
CA1sr: Hippocampal formation, Ammon’s horn, Field CA1, stratum radiatum
CA1sp: Hippocampal formation, Ammon’s horn, Field CA1, pyramidal layer
CA2: Hippocampal formation, Ammon’s horn, Field CA2
CA2sp: Hippocampal formation, Ammon’s horn, Field CA2, pyramidal layer
CA2so: Hippocampal formation, Ammon’s horn, Field CA2, stratum oriens
CA1sr: Hippocampal formation, Ammon’s horn, Field CA2, stratum radiatum
CA2: Hippocampal formation, Ammon’s horn, Field CA3
CA3sp: Hippocampal formation, Ammon’s horn, Field CA3, pyramidal layer
CA3sp: Hippocampal formation, Ammon’s horn, Field CA3, pyramidal layer
CA3so: Hippocampal formation, Ammon’s horn, Field CA3, stratum oriens
CA3sr: Hippocampal formation, Ammon’s horn, Field CA3, stratum radiatum
CA3slu: Hippocampal formation, Ammon’s horn, Field CA3, stratum lucidum
DG: Hippocampal formation, dentate gyrus
DG-mo: Hippocampal formation, dentate gyrus, molecular layer
DG-sg: Hippocampal formation, dentate gyrus, granule cell layer
SUB: Hippocampal formation, subiculum
ENT: Hippocampal formation, entorhinal area
ENTm: Hippocampal formation, entorhinal area, medial part, dorsal zone
ENTmv: Hippocampal formation, entorhinal area, medial part, ventral zone
ENTl: Hippocampal formation, entorhinal area, lateral part
sAMY: Cerebral nuclei, striatum, striatum-like amygdalar nuclei
PAL: Cerebral nuclei, pallidum
PALv: Cerebral nuclei, pallidum, ventral region
PALd: Cerebral nuclei, pallidum, dorsal region
BST: Cerebral nuclei, pallidum, caudal region, bed nuclei of the stria terminalis
MSC: Cerebral nuclei, pallidum, medial region, medial septal complex
STRv: Cerebral nuclei, striatum ventral region
OT: Cerebral nuclei, striatum ventral region, olfactory tubercle
FS: Cerebral nuclei, striatum ventral region, fundus of striatum
ACB: Cerebral nuclei, striatum ventral region, nucleus accumbens
CP: Cerebral nuclei, striatum dorsal region, caudoputamen
LSX: Cerebral nuclei, striatum dorsal region, lateral septal complex
CB: Cerebellum
DN: Cerebellum, dentate nucleus
IP: Cerebellum, interposed nucleus
FN: Cerebellum, fastigial nucleus
SIM: Cerebellum, hemispheric regions, simple lobule
ANcr1: Cerebellum, hemispheric regions, ansiform lobule, crus 1
ANcr2: Cerebellum, hemispheric regions, ansiform lobule, crus 2
PRM: Cerebellum, hemispheric regions, paramedian lobule
COPY: Cerebellum, hemispheric regions, copula pyramidis
FL: Cerebellum, hemispheric regions, flocculus
PFL: Cerebellum, hemispheric regions, paraflocculus
LING: Cerebellum, vermal regions, lingula (I)
CUL: Cerebellum, vermal regions, culmen
DEC: Cerebellum, vermal regions, declive (VI)
PYR: Cerebellum, vermal regions, pyramus (VIII)
NOD: Cerebellum, vermal regions, nodulus (X)
CENT: Cerebellum, vermal regions, central lobule
FOTU: Cerebellum, vermal regions, folium-tuber vermis (VII)
UVU: Cerebellum, vermal regions, uvula (IX)
HY: Brain stem, hypothalamus
MPO: Brain stem, hypothalamus, medial preoptic nucleus
MBO: Brain stem, hypothalamus, mammillary bodies
TH: Brain stem, thalamus
P: Brain stem, hindbrain, pons
SOC: Brain stem, hindbrain, pons, superior olivary complex
PRNr: Brain stem, hindbrain, pons, pontine nucleus
MY: Brain stem, hindbrain, medulla
IO: Brain stem, hindbrain, medulla, inferior olivary complex
CU: Brain stem, hindbrain, medulla, cuneate nucleus
MB: Brain stem, midbrain
SCs: Brain stem, midbrain, colliculus: superior
IC: Brain stem, midbrain, colliculus: inferior
IPN: Brain stem, midbrain, interpendunclar nucleus
PAG: Brain stem, midbrain, periaqueductal grey
OLF: Olfactory areas
MOB: Olfactory area, main olfactory bulb
MOBgl: Olfactory area, main olfactory bulb, glomerular layer
MOBopl: Olfactory area, main olfactory bulb, outer plexiform layer
MOBmi: Olfactory area, main olfactory bulb, mitral layer
MOBipl: Olfactory area, main olfactory bulb, inner plexiform layer
MOBgr: Olfactory area, main olfactory bulb, granule layer
AOB: Olfactory area, accessory olfactory bulb
AOBgl: Olfactory area, accessory olfactory bulb, glomerular layer
AOBgr: Olfactory area, accessory olfactory bulb, glomerular layer
AON: Olfactory area, anterior olfactory nucleus
PIR: Olfactory area, piriform area
TT: Olfactory area, taenia tecta
TTd: Olfactory area, taenia tecta, dorsal part
TTv: Olfactory area, taenia tecta, ventral part
COA: Olfactory area, cortical amygdalar area
COApl: Olfactory area, cortical amygdalar area, posterior part, lateral zone
COApm: Olfactory area, cortical amygdalar area, posterior part, medial zone
PAA: Olfactory area, piriform-amygdalar area
mfbc: Fiber tracts, medial forebrain bundle system, cerebrum related
lfbs: Fiber tracts, lateral forebrain bundle system
cm: Fiber tracts, cranial nerves
mfsbshy: Fiber tracts, medial forebrain bundle system, hypothalamus related
cbf: Fiber tracts, cerebellum related fiber tracts
arb: Fiber tracts, cerebellum related fiber tracts, arbor vitae
VL: Ventricular systems, lateral ventricle
V4: Ventricular systems, fourth ventricle
AQ: Ventricular systems, cerebral aqueduct
V3: Ventricular systems, third ventricle
SEZ: Ventricular systems, lateral ventricle, subependymal zone

## References

1. Dominguez-Bello MG, Costello EK, Contreras M, Magris M, Hidalgo G, Fierer N, et al. Delivery mode shapes the acquisition and structure of the initial microbiota across multiple body habitats in newborns. Proc Natl Acad Sci USA. 2010 Jun 29;107(26):11971–5.

2. Grölund MM, Lehtonen OP, Eerola E, Kero P. Fecal Microflora in Healthy Infants Born by Different Methods of Delivery: Permanent Changes in Intestinal Flora After Cesarean Delivery: Journal of Pediatric Gastroenterology & Nutrition. 1999 Jan;28(1):19–25.

3. Henrick BM, Rodriguez L, Lakshmikanth T, Pou C, Henckel E, Arzoomand A, et al. Bifidobacteria-mediated immune system imprinting early in life. Cell. 2021 Jul;184(15):3884-3898.e11.

4. Wu HJ, Wu E. The role of gut microbiota in immune homeostasis and autoimmunity. Gut Microbes. 2012 Jan;3(1):4–14.

5. Martin AM, Sun EW, Rogers GB, Keating DJ. The Influence of the Gut Microbiome on Host Metabolism Through the Regulation of Gut Hormone Release. Front Physiol. 2019 Apr 16;10:428.

6. Agus A, Clément K, Sokol H. Gut microbiota-derived metabolites as central regulators in metabolic disorders. Gut. 2021 Jun;70(6):1174–82.

7. Rhys-Jones D, Climie RE, Gill PA, Jama HA, Head GA, Gibson PR, et al. Microbial Interventions to Control and Reduce Blood Pressure in Australia (MICRoBIA): rationale and design of a double-blinded randomised cross-over placebo controlled trial. Trials. 2021 Dec;22(1):496.

8. Matenchuk BA, Mandhane PJ, Kozyrskyj AL. Sleep, circadian rhythm, and gut microbiota. Sleep Medicine Reviews. 2020 Oct;53:101340.

9. Breit S, Kupferberg A, Rogler G, Hasler G. Vagus Nerve as Modulator of the Brain–Gut Axis in Psychiatric and Inflammatory Disorders. Front Psychiatry. 2018 Mar 13;9:44.

10. Cryan JF, O’Riordan KJ, Cowan CSM, Sandhu KV, Bastiaanssen TFS, Boehme M, et al. The Microbiota-Gut-Brain Axis. Physiological Reviews. 2019 Oct 1;99(4):1877–2013.

11. Gasparrini AJ, Wang B, Sun X, Kennedy EA, Hernandez-Leyva A, Ndao IM, et al. Persistent metagenomic signatures of early-life hospitalization and antibiotic treatment in the infant gut microbiota and resistome. Nat Microbiol. 2019 Sep 9;4(12):2285–97.

12. Bokulich NA, Chung J, Battaglia T, Henderson N, Jay M, Li H, et al. Antibiotics, birth mode, and diet shape microbiome maturation during early life. Sci Transl Med [Internet]. 2016 Jun 15 [cited 2022 Dec 19];8(343). Available from: https://www.science.org/doi/10.1126/scitranslmed.aad7121

13. Slob EMA, Brew BK, Vijverberg SJH, Dijs T, van Beijsterveldt CEM, Koppelman GH, et al. Early-life antibiotic use and risk of attention-deficit hyperactivity disorder and autism spectrum disorder: results of a discordant twin study. International Journal of Epidemiology. 2021 May 17;50(2):475– 84.

14. Slykerman RF, Thompson J, Waldie KE, Murphy R, Wall C, Mitchell EA. Antibiotics in the first year of life and subsequent neurocognitive outcomes. Acta Paediatr. 2017 Jan;106(1):87–94.

15. Lu J, Lu L, Yu Y, Cluette-Brown J, Martin CR, Claud EC. Effects of Intestinal Microbiota on Brain Development in Humanized Gnotobiotic Mice. Sci Rep. 2018 Dec;8(1):5443.

16. Slykerman RF, Coomarasamy C, Wickens K, Thompson JMD, Stanley TV, Barthow C, et al. Exposure to antibiotics in the first 24 months of life and neurocognitive outcomes at 11 years of age. Psychopharmacology. 2019 May;236(5):1573–82.

17. Valles-Colomer M, Falony G, Darzi Y, Tigchelaar EF, Wang J, Tito RY, et al. The neuroactive potential of the human gut microbiota in quality of life and depression. Nat Microbiol. 2019 Apr;4(4):623–32.

18. Kratsman N, Getselter D, Elliott E. Sodium butyrate attenuates social behavior deficits and modifies the transcription of inhibitory/excitatory genes in the frontal cortex of an autism model. Neuropharmacology. 2016 Mar;102:136–45.

19. Kundu P, Lee HU, Garcia-Perez I, Tay EXY, Kim H, Faylon LE, et al. Neurogenesis and prolongevity signaling in young germ-free mice transplanted with the gut microbiota of old mice. Sci Transl Med. 2019 Nov 13;11(518):eaau4760.

20. Wei GZ, Martin KA, Xing PY, Agrawal R, Whiley L, Wood TK, et al. Tryptophan-metabolizing gut microbes regulate adult neurogenesis via the aryl hydrocarbon receptor. Proc Natl Acad Sci USA. 2021 Jul 6;118(27):e2021091118.

21. Vuong HE, Pronovost GN, Williams DW, Coley EJL, Siegler EL, Qiu A, et al. The maternal microbiome modulates fetal neurodevelopment in mice. Nature. 2020 Oct 8;586(7828):281–6.

22. Heijtz RD, Wang S, Anuar F, Qian Y, Björkholm B, Samuelsson A, et al. Normal gut microbiota modulates brain development and behavior. Proc Natl Acad Sci USA. 2011 Feb 15;108(7):3047– 52.

23. Neufeld KM, Kang N, Bienenstock J, Foster JA. Reduced anxiety-like behavior and central neurochemical change in germ-free mice: Behavior in germ-free mice. Neurogastroenterology & Motility. 2011 Mar;23(3):255–e119.

24. Agranyoni O, Meninger-Mordechay S, Uzan A, Ziv O, Salmon-Divon M, Rodin D, et al. Gut microbiota determines the social behavior of mice and induces metabolic and inflammatory changes in their adipose tissue. npj Biofilms Microbiomes. 2021 Mar 19;7(1):28.

25. Desbonnet L, Clarke G, Shanahan F, Dinan TG, Cryan JF. Microbiota is essential for social development in the mouse. Mol Psychiatry. 2014 Feb;19(2):146–8.

26. Wu WL, Adame MD, Liou CW, Barlow JT, Lai TT, Sharon G, et al. Microbiota regulate social behaviour via stress response neurons in the brain. Nature. 2021 Jul 15;595(7867):409–14.

27. Aswendt M, Green C, Sadler R, Llovera G, Dzikowski L, Heindl S, et al. The gut microbiota modulates brain network connectivity under physiological conditions and after acute brain ischemia. iScience. 2021 Oct;24(10):103095.

28. Oguz I, Yaxley R, Budin F, Hoogstoel M, Lee J, Maltbie E, et al. Comparison of Magnetic Resonance Imaging in Live vs. Post Mortem Rat Brains. Aoki I, editor. PLoS ONE. 2013 Aug 13;8(8):e71027.

29. Hennig J, Nauerth A, Friedburg H. RARE imaging: A fast imaging method for clinical MR. Magn Reson Med. 1986 Dec;3(6):823–33.

30. Dorr AE, Lerch JP, Spring S, Kabani N, Henkelman RM. High resolution three-dimensional brain atlas using an average magnetic resonance image of 40 adult C57Bl/6J mice. NeuroImage. 2008 Aug;42(1):60–9.

31. Guernier V, Brennan B, Yakob L, Milinovich G, Clements ACA, Soares Magalhaes RJ. Gut microbiota disturbance during helminth infection: can it affect cognition and behaviour of children? BMC Infect Dis. 2017 Dec;17(1):58.

32. Tamana SK, Tun HM, Konya T, Chari RS, Field CJ, Guttman DS, et al. Bacteroides-dominant gut microbiome of late infancy is associated with enhanced neurodevelopment. Gut Microbes. 2021 Jan 1;13(1):1930875.

33. El Aidy S, Derrien M, Merrifield CA, Levenez F, Doré J, Boekschoten MV, et al. Gut bacteria– host metabolic interplay during conventionalisation of the mouse germfree colon. ISME J. 2013 Apr;7(4):743–55.

34. Chevalier G, Siopi E, Guenin-Macé L, Pascal M, Laval T, Rifflet A, et al. Effect of gut microbiota on depressive-like behaviors in mice is mediated by the endocannabinoid system. Nat Commun. 2020 Dec;11(1):6363.

35. Abdelli LS, Samsam A, Naser SA. Propionic Acid Induces Gliosis and Neuro-inflammation through Modulation of PTEN/AKT Pathway in Autism Spectrum Disorder. Sci Rep. 2019 Jun 19;9(1):8824.

36. Erny D, Hrabě de Angelis Al, Jaitin D, Wieghofer P, Staszewski O, David E, et al. Host microbiota constantly control maturation and function of microglia in the CNS. Nat Neurosci. 2015 Jul;18(7):965–77.

37. Goodrich JK, Waters JL, Poole AC, Sutter JL, Koren O, Blekhman R, et al. Human Genetics Shape the Gut Microbiome. Cell. 2014 Nov;159(4):789–99.

38. Rosenfeld CS. Gut Dysbiosis in Animals Due to Environmental Chemical Exposures. Front Cell Infect Microbiol. 2017 Sep 8;7:396.

39. Ramirez J, Guarner F, Bustos Fernandez L, Maruy A, Sdepanian VL, Cohen H. Antibiotics as Major Disruptors of Gut Microbiota. Front Cell Infect Microbiol. 2020 Nov 24;10:572912.

40. Bai X, Narayanan A, Nowak P, Ray S, Neogi U, Sönnerborg A. Whole-Genome Metagenomic Analysis of the Gut Microbiome in HIV-1-Infected Individuals on Antiretroviral Therapy. Front Microbiol. 2021 Jun 25;12:667718.

41. Kim CY, Lee M, Yang S, Kim K, Yong D, Kim HR, et al. Human reference gut microbiome catalog including newly assembled genomes from under-represented Asian metagenomes. Genome Med. 2021 Dec;13(1):134.

42. Li F, Yang S, Zhang L, Qiao L, Wang L, He S, et al. Comparative metagenomics analysis reveals how the diet shapes the gut microbiota in several small mammals. Ecology and Evolution [Internet]. 2022 Jan [cited 2022 Jun 7];12(1). Available from: https://onlinelibrary.wiley.com/doi/10.1002/ece3.8470

43. Kieser S, Zdobnov EM, Trajkovski M. Comprehensive mouse microbiota genome catalog reveals major difference to its human counterpart. Nagarajan N, editor. PLoS Comput Biol. 2022 Mar 8;18(3):e1009947.

44. Kennedy EA, King KY, Baldridge MT. Mouse Microbiota Models: Comparing Germ-Free Mice and Antibiotics Treatment as Tools for Modifying Gut Bacteria. Front Physiol. 2018 Oct 31;9:1534.

45. Park JC, Im SH. Of men in mice: the development and application of a humanized gnotobiotic mouse model for microbiome therapeutics. Exp Mol Med. 2020 Sep;52(9):1383–96.

46. Kundu P, Blacher E, Elinav E, Pettersson S. Our Gut Microbiome: The Evolving Inner Self. Cell. 2017 Dec;171(7):1481–93.

47. Choe M s., Ortiz-Mantilla S, Makris N, Gregas M, Bacic J, Haehn D, et al. Regional Infant Brain Development: An MRI-Based Morphometric Analysis in 3 to 13 Month Olds. Cerebral Cortex. 2013 Sep 1;23(9):2100–17.

48. Cheong JLY, Thompson DK, Spittle AJ, Potter CR, Walsh JM, Burnett AC, et al. Brain Volumes at Term-Equivalent Age Are Associated with 2-Year Neurodevelopment in Moderate and Late Preterm Children. The Journal of Pediatrics. 2016 Jul;174:91-97.e1.

49. Gómez LJ, Dooley JC, Sokoloff G, Blumberg MS. Parallel and Serial Sensory Processing in Developing Primary Somatosensory and Motor Cortex. J Neurosci. 2021 Apr 14;41(15):3418–31.

50. Laurienti PJ, Wallace MT, Maldjian JA, Susi CM, Stein BE, Burdette JH. Cross-modal sensory processing in the anterior cingulate and medial prefrontal cortices. Hum Brain Mapp. 2003 Aug;19(4):213–23.

51. Xerri C, Zennou-Azogui Y. Interplay between Primary Cortical Areas and Crossmodal Plasticity. In: Heinbockel T, Zhou Y, editors. Connectivity and Functional Specialization in the Brain [Internet]. IntechOpen; 2021 [cited 2022 Aug 28]. Available from: https://www.intechopen.com/books/connectivity-and-functional-specialization-in-the-brain/interplay-between-primary-cortical-areas-and-crossmodal-plasticity

52. Medinaceli Quintela R, Brunert D, Rothermel M. Functional role of the anterior olfactory nucleus in sensory information processing. Neuroforum. 2022 Aug 26;28(3):169–75.

53. Feigin L, Tasaka G, Maor I, Mizrahi A. Sparse Coding in Temporal Association Cortex Improves Complex Sound Discriminability. J Neurosci. 2021 Aug 18;41(33):7048–64.

54. Dash S, Clarke G, Berk M, Jacka FN. The gut microbiome and diet in psychiatry: focus on depression. Current Opinion in Psychiatry. 2015 Jan;28(1):1–6.

55. Habata K, Cheong Y, Kamiya T, Shiotsu D, Omori IM, Okazawa H, et al. Relationship between sensory characteristics and cortical thickness/volume in autism spectrum disorders. Transl Psychiatry. 2021 Dec;11(1):616.

56. Tayler KK, Tanaka KZ, Reijmers LG, Wiltgen BJ. Reactivation of Neural Ensembles during the Retrieval of Recent and Remote Memory. Current Biology. 2013 Jan;23(2):99–106.

57. Jinks AL, McGregor IS. Modulation of anxiety-related behaviours following lesions of the prelimbic or infralimbic cortex in the rat. Brain Research. 1997 Oct;772(1–2):181–90.

58. Schulz-Klaus B, Fendt M, Schnitzler HU. Temporary inactivation of the rostral perirhinal cortex induces an anxiolytic-like effect on the elevated plus-maze and on the yohimbine-enhanced startle response. Behavioural Brain Research. 2005 Sep;163(2):168–73.

59. Cádiz-Moretti B, Abellán-Álvaro M, Pardo-Bellver C, Martínez-García F, Lanuza E. Afferent and Efferent Connections of the Cortex-Amygdala Transition Zone in Mice. Front Neuroanat [Internet]. 2016 Dec 23 [cited 2022 Jun 7];10. Available from: http://journal.frontiersin.org/article/10.3389/fnana.2016.00125/full

60. Liu HX, Rocha CS, Dandekar S, Wan YJY. Functional analysis of the relationship between intestinal microbiota and the expression of hepatic genes and pathways during the course of liver regeneration. J Hepatol. 2016 Mar;64(3):641–50.

61. Levinson M, Kolenda JP, Alexandrou GJ, Escanilla O, Cleland TA, Smith DM, et al. Context-dependent odor learning requires the anterior olfactory nucleus. Behavioral Neuroscience. 2020 Aug;134(4):332–43.

62. Xu M, Minagawa Y, Kumazaki H, Okada K ichi, Naoi N. Prefrontal Responses to Odors in Individuals With Autism Spectrum Disorders: Functional NIRS Measurement Combined With a Fragrance Pulse Ejection System. Front Hum Neurosci. 2020 Oct 8;14:523456.

63. Koehler L, Fournel A, Albertowski K, Roessner V, Gerber J, Hummel C, et al. Impaired Odor Perception in Autism Spectrum Disorder Is Associated with Decreased Activity in Olfactory Cortex. Chemical Senses. 2018 Sep 22;43(8):627–34.

64. Deems DA, Doty RL, Settle RG, Moore-Gillon V, Shaman P, Mester AF, et al. Smell and Taste Disorders, A Study of 750 Patients From the University of Pennsylvania Smell and Taste Center. Archives of Otolaryngology - Head and Neck Surgery. 1991 May 1;117(5):519–28.

65. Moberg P. Olfactory Dysfunction in Schizophrenia A Qualitative and Quantitative Review. Neuropsychopharmacology. 1999 Sep;21(3):325–40.

66. Balmer CW, LaMantia AS. Noses and neurons: Induction, morphogenesis, and neuronal differentiation in the peripheral olfactory pathway. Dev Dyn. 2005 Nov;234(3):464–81.

67. Feinstein P, Bozza T, Rodriguez I, Vassalli A, Mombaerts P. Axon Guidance of Mouse Olfactory Sensory Neurons by Odorant Receptors and the β2 Adrenergic Receptor. Cell. 2004 Jun;117(6):833–46.

68. Pluznick JL, Protzko RJ, Gevorgyan H, Peterlin Z, Sipos A, Han J, et al. Olfactory receptor responding to gut microbiota-derived signals plays a role in renin secretion and blood pressure regulation. Proc Natl Acad Sci USA. 2013 Mar 12;110(11):4410–5.

69. Maraci Ö, Engel K, Caspers BA. Olfactory Communication via Microbiota: What Is Known in Birds? Genes. 2018 Jul 31;9(8):387.

70. Dash S, Syed YA, Khan MR. Understanding the Role of the Gut Microbiome in Brain Development and Its Association With Neurodevelopmental Psychiatric Disorders. Front Cell Dev Biol. 2022 Apr 14;10:880544.

71. Glebov K, Löchner M, Jabs R, Lau T, Merkel O, Schloss P, et al. Serotonin stimulates secretion of exosomes from microglia cells: Serotonin Stimulates Microglial Exosome Release. Glia. 2015 Apr;63(4):626–34.

72. Feng W, Wang Y, Liu ZQ, Zhang X, Han R, Miao YZ, et al. Microglia activation contributes to quinolinic acid-induced neuronal excitotoxicity through TNF-α. Apoptosis. 2017 May;22(5):696– 709.

73. Stoeckli ET. Morphogens and Neural Development. In: Binder MD, Hirokawa N, Windhorst U, editors. Encyclopedia of Neuroscience [Internet]. Berlin, Heidelberg: Springer Berlin Heidelberg; 2009 [cited 2022 Jun 7]. p. 2397–401. Available from: http://link.springer.com/10.1007/978-3-540-29678-2_3562

74. Semple BD, Blomgren K, Gimlin K, Ferriero DM, Noble-Haeusslein LJ. Brain development in rodents and humans: Identifying benchmarks of maturation and vulnerability to injury across species. Progress in Neurobiology. 2013 Jul;106–107:1–16.

75. Aldridge K, Cole KK, Moffitt Gunn AJ, Peck D, White DA, Christ SE. The effects of early-treated phenylketonuria on volumetric measures of the cerebellum. Molecular Genetics and Metabolism Reports. 2020 Dec;25:100647.

76. Thanos PK, Michaelides M, Gispert JD, Pascau J, Soto-Montenegro ML, Desco M, et al. Differences in response to food stimuli in a rat model of obesity: in-vivo assessment of brain glucose metabolism. Int J Obes. 2008 Jul;32(7):1171–9.

77. Tang W, Meng Z, Li N, Liu Y, Li L, Chen D, et al. Roles of Gut Microbiota in the Regulation of Hippocampal Plasticity, Inflammation, and Hippocampus-Dependent Behaviors. Front Cell Infect Microbiol. 2021 Jan 27;10:611014.

78. Dieleman N, Koek HL, Hendrikse J. Short-term mechanisms influencing volumetric brain dynamics. NeuroImage: Clinical. 2017;16:507–13.

79. Bäckhed F, Ding H, Wang T, Hooper LV, Koh GY, Nagy A, et al. The gut microbiota as an environmental factor that regulates fat storage. Proc Natl Acad Sci USA. 2004 Nov 2;101(44):15718–23.

